# Demographic inference under the coalescent in a spatial continuum

**DOI:** 10.1101/042135

**Authors:** Stéphane Guindon, Hongbin Guo, David Welch

**Author notes:** Corresponding author: Stéphane Guindon. Current address: LIRMM, Bâtiment 5 - 860 rue de St Priest 34095 Montpellier cedex 5. Phone: +33/0 467 14 97 00.

## Abstract

Understanding population dynamics from the analysis of molecular and spatial data requires sound statistical modeling. Current approaches assume that populations are naturally partitioned into discrete demes, thereby failing to be relevant in cases where individuals are scattered on a spatial continuum. Other models predict the formation of increasingly tight clusters of individuals in space, which, again, conflicts with biological evidence. Building on recent theoretical work, we introduce a new genealogy-based inference framework that alleviates these issues. This approach effectively implements a stochastic model in which the distribution of individuals is homogeneous and stationary, thereby providing a relevant null model for the fluctuation of genetic diversity in time and space. Importantly, the spatial density of individuals in a population and their range of dispersal during the course of evolution are two parameters that can be inferred separately with this method. The validity of the new inference framework is confirmed with extensive simulations and the analysis of influenza sequences collected over five seasons in the USA.

## 1 Introduction

Kingman’s coalescent [24] is a cornerstone of population genetics. It provides a mathematical framework in which the effective size of a population can be estimated through the comparative analysis of genetic data from a sample of individuals. The simplicity and utility of the coalescent explains its popularity in biology (see [32] for a review). In its simplest form, the coalescent defines the probability density of a genealogy of individuals sampled from a constant size, panmictic population. However, the panmixia assumption becomes problematic when considering the spatial distribution of individuals as degree of kinship is generally correlated with geographic distance [37, 34, 25].

The coalescent was thus extended to incorporate spatial information. Under the so-called structured coalescent [21, 33], the population is partitioned into demes, each deme corresponding to a geographic entity. Sub-populations within each deme are panmictic and only individuals in the same deme can coalesce. Migrations between demes are governed by an homogeneous Markov process with the migration rate assumed to be small and estimated from the combination of spatial and genetic data. The structured coalescent has obvious connections with standard models in population genetics, namely the island model [52, 30] and the stepping stone models [29, 23] for which mathematical properties are well understood. Inference under the structured coalescent using maximum-likelihood [7, 8] and Bayesian techniques [15, 6, 46] has led to important advances in biology [e.g., 40], but is limited for computational reasons to a relatively small number of demes (typically less than ten) which are assumed known a priori [46].

But many natural populations are not subdivided into discrete demes. Instead, they display a gradient of kinship across a continuous landscape. In seminal works, Wright [53] and Malécot [28] proposed an extension of the Wright-Fisher model that incorporates continuous spatial information. Under the so-called “isolation-by-distance” (IBD) model, individuals are uniformly distributed on the landscape and the locations of offspring are random draws from a Normal distribution with mean given by the parental position. Mathematical expressions were derived for the probability that two alleles are identical by descent as a function of their spatial distance. However, Felsenstein [16] showed that some of the assumptions of the IBD model are inconsistent. A population evolving under this process displays ‘clumping’ of individuals, which contradicts the uniformity assumption. The IBD model may provide a suitable inference framework when considering short time scales over which clumping can safely be ignored. In the general case, however, it is preferable that the spatial distribution of the population is described by a stationary process.

Sawyer and Felsenstein [43] addressed this issue in a modified version of the IBD model where the spatial distribution of individuals is governed by a Poisson random field with density constant in time. However, their model relies on the assumption that each pair of parents produces exactly two offspring, which is constraining from a biological perspective. Moreover, their approach applies to the special case of a one-dimensional habitat and generalization to two dimensions leads to mathematical difficulties [31].

Wilkins and Wakeley [51] proposed a different approach in which a population is uniformly distributed along a one-dimensional finite habitat with the location of parents correlated to that of offspring. Importantly, each individual occupies an interval inversely proportional to the size of the population, thereby ensuring the population density is regulated at all points in space and time. This model assumes that the spatial position of each lineage is subject to a diffusion process backward in time, with the habitat having reflecting boundaries. The authors were able to derive analytic formula for the distribution of the time to coalescence of a pair of sampled lineages. Wilkins [50] later proposed a generalization of this model to two-dimensional landscapes. However, Barton, Etheridge and Véber (2010b) suggested that this approach is sampling inconsistent, i.e., the distribution of the time to coalescence of a pair of sampled lineages depends on the size of the sample under consideration. Estimates of parameters of this model may thus be difficult to interpret in practice.

More recently, Lemey et al. (2009) proposed a model whereby spatial location is considered as a discrete character evolving along lineages according to a continuous-time Markov chain. Unfortunately, this approach suffers serious limitations. First, estimates of rates of migration are influenced by spatial variations in sampling intensity. Moreover, non-uniformity of population density is ignored when calculating the density of the genealogy. Also, this model is a discrete approximation of the IBD model when the migration process is isotropic and thus suffers from the same shortcomings. Altogether, while this approach is efficient from a computational perspective, it provides biased estimates of demographic parameters in particular simulation settings [13] and should thus be used with great caution.

In a recent series of articles [14, 9, 3, 5, 47, 4], Barton, Etheridge and colleagues described a new process, called the spatial λ-Fleming-Viot process, for studying the evolution of populations on a continuous landscape. Malécot’s approach and related models consider that the time of death and reproduction of individuals are governed by a random process running along every lineage in the evolving population. The new model assumes instead that the time and position of these events are independent of the spatial location of lineages. The authors describe the forward-in-time dynamics of a population evolving on an unbounded spatial continuum, and the corresponding backward-in-time process that characterizes the genealogy of sampled individuals.

The mathematical properties of the spatial λ-Fleming-Viot model have been studied extensively [9, 3, 47, 4]. In particular, it has been shown that this model does not suffer from sampling inconsistency or clumping issues. Barton et al. [2] showed that the analysis of pairs of sub-populations provides information about neighborhood size, i.e., the product of the effective population density by the dispersal intensity. These last two quantities are relevant from a biological perspective and, ideally, one would like to estimate each of them separately instead of their product.

In this study, we perform Bayesian inference under the spatial Λ-Fleming-Viot model applied to multiple individuals taken jointly. Using extensive simulations, we demonstrate the accuracy and precision with which the parameters of this model can be estimated. We compare our estimates to that obtained using two popular inference techniques, i.e., the regression on fixation index (Fst) values [41] and the structured coalescent [21, 33]. Our results demonstrate the good performance of our approach in these conditions. They also indicate that the proposed framework permits the estimation of population density and dispersal intensity as two separate (i.e., identifiable) parameters, thereby going beyond pairwise analyses. We further illustrate the validity of this new technique through the analysis of H1N1 sequences collected over five flu seasons in the USA. We show that the 2009 flu pandemic had distinct population dynamics compared to more recent seasons with smaller than usual neighborhood size and larger than usual dispersal distance.

## 2 The model

The spatial Λ-Fleming-Viot model (noted as ΛV from here on for the sake of brevity) assumes that reproduction, dispersal and death of lineages result from ‘events’ that occur at locations independent from that of individuals forming the population under scrutiny. In the following, we refer to these events as reproduction/extinction or REX events. From a biological perspective, one REX corresponds to either (i) a single reproduction event accompanied by extinction of the parent with the offspring dispersing over long distances or (ii) a sum of multiple reproduction and extinction events, each reproduction accompanied by dispersal of the offspring over short distances. In any case, the average time between two successive REX in a given lineage is proportional to the generation time of the species under scrutiny.

### 2-a Forward-in-time dynamics of the population

We assume that a population inhabits a finite habitat represented by a rectangle *R*(*h, ω*), with height *h* and width *ω* known *a priori*. Migrations crossing the boundaries of the rectangle are forbidden. This differs slightly from Barton, Etheridge and colleagues who assume that the population is distributed on a two-dimensional torus or on ℝ^2^.

Initial locations of individuals are determined by a homogeneous Poisson point process with intensity *ρ*, producing a uniform distribution over *R*(*h, ω*). REXs occur at exponentially distributed waiting times with rate parameter λ. The center of each REX is chosen uniformly at random across the habitat. Each lineage then dies with some probability which depends on its distance to the center. New lineages are born at an intensity that also depends on the distance to the center. The choice of kernel modeling the dependency of the rates of birth and death as a function of distance from the center is flexible [see 4]. We choose a Normal kernel.

Let *c*_*i*_ := (*x*_*i*_, *y*_*i*_) be the center of the REX occurring at time *t*_*i*_. Suppose lineage *k* has location 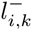 at time 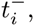 i.e., just before the event occurs (going forward in time). That lineage dies at *t*_*i*_ with probability

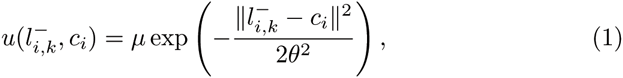

where 0 < *μ* ≤ 1. We will refer to *μ* and *θ* as the death and radius parameters from here on.

Individuals are born at time *t*_*i*_ and location *l* according to an inhomogeneous Poisson point process with intensity *ρu*(*l,c*_*i*_)*dl*. Therefore, the expected size of the population does not change after each REX event and the spatial distribution of the population is still uniform, making this process stationary.

Finally, all individuals born at *t*_*i*_ share a common parent who is chosen from among the individuals existing immediately before the REX event. The probability that lineage *k* with location 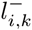 is selected as parent is then

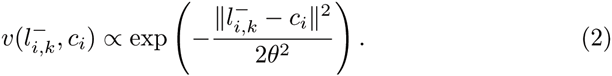

### 2-b Backward-in-time dynamics of a sample

When considering a sample from the present population (corresponding to time *t*_0_ = 0), its ancestry is traced towards the past (corresponding to *t* < 0) as follows. REXs occur at the same rate λ as in the forward-in-time process and centers (i.e., values of *c*_*i*_) are still uniform on *R*(*h, ω*). Suppose a REX event takes place at time *t*_*i*_ and that lineage *k* has position 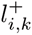 at time 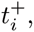 i.e., immediately before the event that took place (going backward in time). *k* changes location at the event, i.e., it is ‘hit’ by the REX event, with probability 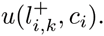. When it is hit, *k* jumps to a new, ancestral, location 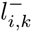 with probability density proportional to 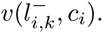. When multiple sampled lineages are hit by the event, all of them coalesce at time *t*_*i*_ and location 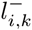.

### 2-c Interpretation of parameters

The ΛV model has three parameters: λ, *μ* and *θ*. λ has a straightforward interpretation: it is the rate of REX events taking place in the whole habitat. It is also common to refer to the rate per unit area λ′ := λ/s where *s* := *ωh* is the total area. Although the value of this parameter is linked to the expected time elapsed between birth and the age of first reproduction, the precise relationship between λ and generation time depends on the biology of the species under scrutiny.

The probability density of the location of a lineage immediately after being hit (going backward in time) given its location before the event is a bivariate normal density with covariance matrix 2*θ*^2^**I**. 2*θ*^2^ is thus half the expected square Euclidean distance (in one dimension) between a parent and one of its immediate offspring.

Also, considering the backward-in-time process here again, the probability that a lineage is hit by a REX is approximated by 2*πθ*^2^*μ/s*. This probability can be interpreted as the ratio between the rate of events that hit lineages and the rate for both types of events. Therefore, given a fixed radius, the higher *μ*, the higher this ratio.

Wright’s neighborhood size (*𝒩*) and dispersal intensity (*σ*^2^) are two standard parameters in population genetics that can be expressed as functions of λ, *μ* and *θ*. Assuming small values of *θ*, the distribution of the location of a lineage at time *t* conditional on its location at time 0 is normal with variance *σ*^2^*t*, where *σ*^2^ := 4*θ*^4^λ′*πμ*. In the limit where λ → ∞ and *θ*→ 0, i.e., the jumps become increasingly frequent and small, the backward-in-time motion of a single lineage is a Brownian process with diffusion parameter *σ*^2^.

Considering again small values of the radius, the probability that any two lineages coalesce in a REX event is simply the probability that both of them are hit, i.e., (2*πθ*;^2^*μ/s*)^2^ =4*π*^2^*θ*^4^*μ*^2^/*s*^2^. The rate of these events is thus4*π*^2^*θ*^4^*μ*^2^λ/*s*^2^. We define this rate as 1/(2*N*_*e*_), where *N*_*e*_ is the effective size of the (diploid) population. Following the definition of the neighborhood size used in the Wright-Malecot model, i.e.,*𝒩* := 4*πN*_*e*_*σ*^2^/*s*, and using the definitions of *σ^2^* and *N*_*e*_ given above, we obtain *𝒩* = 2/*μ*. The neighborhood size, which can be understood as the expected number of individuals participating in reproduction in a disk or radius 2*σ*, is thus inversely proportional to *μ*. More details on the derivations of the results in this section are given in SI 12 and [5].

## 3 Likelihood

We now derive the likelihood of the ΛV model for data consisting of the locations of a sample of present-day individuals and their ancestors. Let *g* be the timed genealogy describing the ancestral relationships of the sampled lineages. Suppose there are *m* REX events between t0 and the time of the most recent common ancestor (MRCA, the root of g) and that the i-th event occurs at time *t*_*i*_ with center *c*_*i*_. Since we are considering the backward-in-time process, *t*_*i*_ > *t*_*i*+1_ for *i* = 0,…, *m* −1. Let 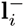 and 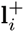 be the vector of locations of the ancestral lineages at time 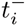 and 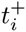 respectively. Also, let 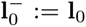 be the known location data. Knowing *g* allows one to determine which locations in 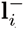 correspond to lineages that were born at *t*_*i*_ and which location in 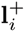 is the parental one.

The likelihood for the observed data l_0_ and imputed data 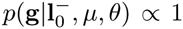, *t*_*i*_ (with *i* =1,… *m*) and *g* is then

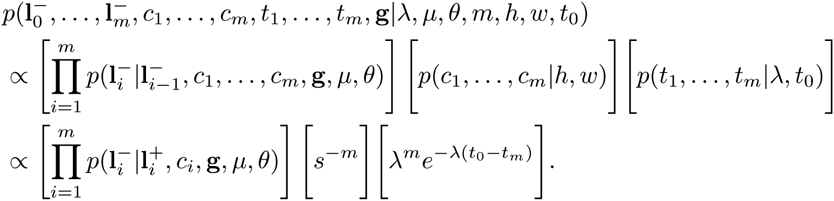

Note that 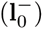, i.e., the locations do not convey any information about the genealogy directly (only genetic sequences do). Let *B*_*i*_ be the set of indices of sampled lineages born at *t*_*i*_. We have:

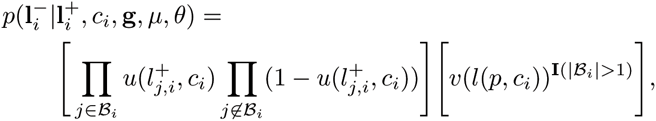

where *l*(*p*, *c*_*i*_) is the location of the parent of all offspring lineages born at *t*_*i*_ and **I**(|*B*_*i*_| > 1) is an indicator function taking value 0 if no offspring were born at REX *i* and 1 otherwise. The likelihood for the observed location data only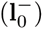 can be obtained via marginalization which we perform via MCMC sampling.

## 4 Simulations

### 4-a Range of parameter values

The habitat is a 10 by 10 square in our simulations. Individuals belonging to the population of interest are never found outside this area. The effective size of the population, *N*_*e*_, was sampled from a uniform distribution on [100, 5000]. Values of the neighborhood size were then obtained by sampling uniformly in [*N*_*e*_ × 10^−3^, *N*_*e*_ × 10^−2^]. Values of *θ* were sampled uniformly in [1.5,4]. When *θ* = 1.5, the probability that the offspring falls at a distance from its parent smaller or equal to 1.0 is approximately 0.5 (the distance is measured here along a single axis). When *θ*; = 4, this probability is approximately 0.25. We thus considered this range of values for *θ* to be broad enough to illustrate medium-and long-range dispersal patterns respectively. The values of *N*_*e*_, *θ* and *𝒩* fully determine that of *μ*, *σ* and λ.

Nucleotide sequences of length 500 bp evolved along the genealogies under the Kimura-2-parameter model [22] with a transition/transversion ratio set to 4.0. The 5% and 95% quantiles of the nucleotide diversity estimated from the sequence alignments hence generated are 0.44% and 1.56% respectively, well in line with that observed in *Drosophila melanogaster* for instance [1].

### 4-b Data collection process

In real experiments, collection of data is rarely uniform over the habitat. Instead, samples are often obtained from disconnected and seemingly randomly scattered regions. In an attempt to mimic these patterns, the sampling scheme used in our simulations relies on throwing random triangles on the habitat. Lineages are then sampled uniformly at random from each of these triangles (see SI 10).

The evolution of a population counting 5,000 individuals was simulated under the forward-in-time process for each simulation. Each of these stopped after 100,000 REXs. These particular values for the population size and number of REXs were selected so that computation could be performed with a reasonable amount of memory. In fact, the size of the population used here is not relevant to the effective population size, which is a function of *λ*, *μ*, *θ* and *s* only.

Sampling sites were then randomly scattered on the landscape, as just explained. A sample of 50 individuals was obtained by selecting individuals uniformly at random within the available sites. In cases where less than 50 individuals happened to be within the sampling regions, new regions and individuals were drawn. This procedure was repeated until the obtention of a sample of size 50. In all simulations, the sampled individuals did coalesce in the time period considered (i.e., the time required to reach 10,000 REXs).

## 5 Bayesian parameter estimation

Samples of genetic sequences, s, along with locations of sampled lineages, l0, are used to infer the parameters of interest, namely λ, *μ* and *θ*. The model also involves multiple nuisance parameters which we impute; the genealogy of the genetic sequences under study and the spatial coordinates of ancestral lineages being the main ones. Random draws from the joint posterior distribution of all these parameters are obtained using MCMC techniques. The posterior density is as follows

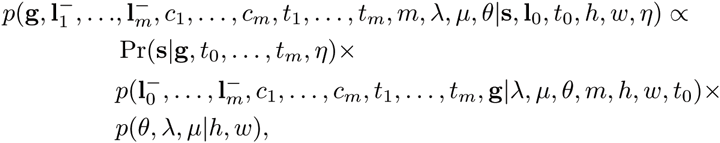

where *η* is the substitution rate. In what follows, we assume that this rate does not vary across lineages nor sites of the alignment and its value is known *a priori*. The parameters (*t*_0_,… ,*t*_*m*_, *η*) fully specify the edge lengths in g in terms of expected number of substitutions (between nucleotides, amino-acids or codons) per position along a sequence. The probability Pr(s|g,*t*_0_, …, *t*_*m*_, *η*) is calculated using Felsenstein’s (1981) pruning algorithm. The joint prior density *p*(*θ*, λ, *μ|h, ω*) is given by the product of the three densities *p*(*θ|h, ω*), *p*(λ|*h, ω*) and *p*(*μ*|*h, ω*). The prior distributions of *μ*, λ and *θ* are uniform on [0,1], [10^−6^,10^+2^] and [0,5] respectively.

Random draws from the joint posterior distribution were obtained using the Metropolis-Hastings algorithm. A total of fifteen operators updating the values of every model parameter, including the tree topology, were implemented (see SI 11). We validated the implementation and verified the correctness of the data simulation algorithm by comparing the distribution of summary statistics in simulated data to that inferred using our sampling technique (see SI 13). For each (real and simulated) data set, the chain was sampled for a maximum of 100 hours on a computer server equipped with 2.7-2.8 GHz CPUs. The sampling halted when the effective sample sizes of λ, *μ* and *σ*^2^ all exceeded 100.

## 6 Results

### 6-a Population density

Neighborhood size (*𝒩*), a quantity proportional to the product of population density and dispersal intensity, can be inferred from pairs of sequences and their spatial coordinates. Indeed, estimates of *𝒩* are often derived from the slope of the regression of Fst values for pairs of sequences on the corresponding geographic distances (see [41] and SI 14). In our simulations, Pearson’s correlation between true neighborhood values and estimates obtained using this technique are equal to 0.074 and −0.006 for two and ten sampling regions respectively. For the ΛV model, estimates of neighborhood sizes are taken as the posterior medians. The correlation between true and estimated values is equal to 0.631 and 0.669 for two and ten sites respectively. A more detailed analysis of the posterior distributions estimated from the simulated data indicates that accurate and precise inference of this parameter is achievable using our technique, at least for values of *𝒩* smaller than ~20 (see SI 15).

The structured coalescent is often used to estimate the effective size of populations in a context similar to that of our simulations: each deme corresponds to a sampling region as opposed to a genuine element of a partitioned population. We used MultiTypeTree [46] from the BEAST2 package [10] to estimate the parameters of the structured coalescent model applied to our simulated data (see SI 17). Posterior medians and 95% credibility intervals for the population densities estimated with the structured coalescent and the ΛV models are presented in Figure 1. The structured coalescent generally overestimates population densities with the strongest biases observed with two demes and more accurate estimates obtained with ten. Estimates obtained under the ΛV model are better overall, both in terms of accuracy and, to a lesser extent, precision, with very little difference between two and ten demes.

**Figure 1:**
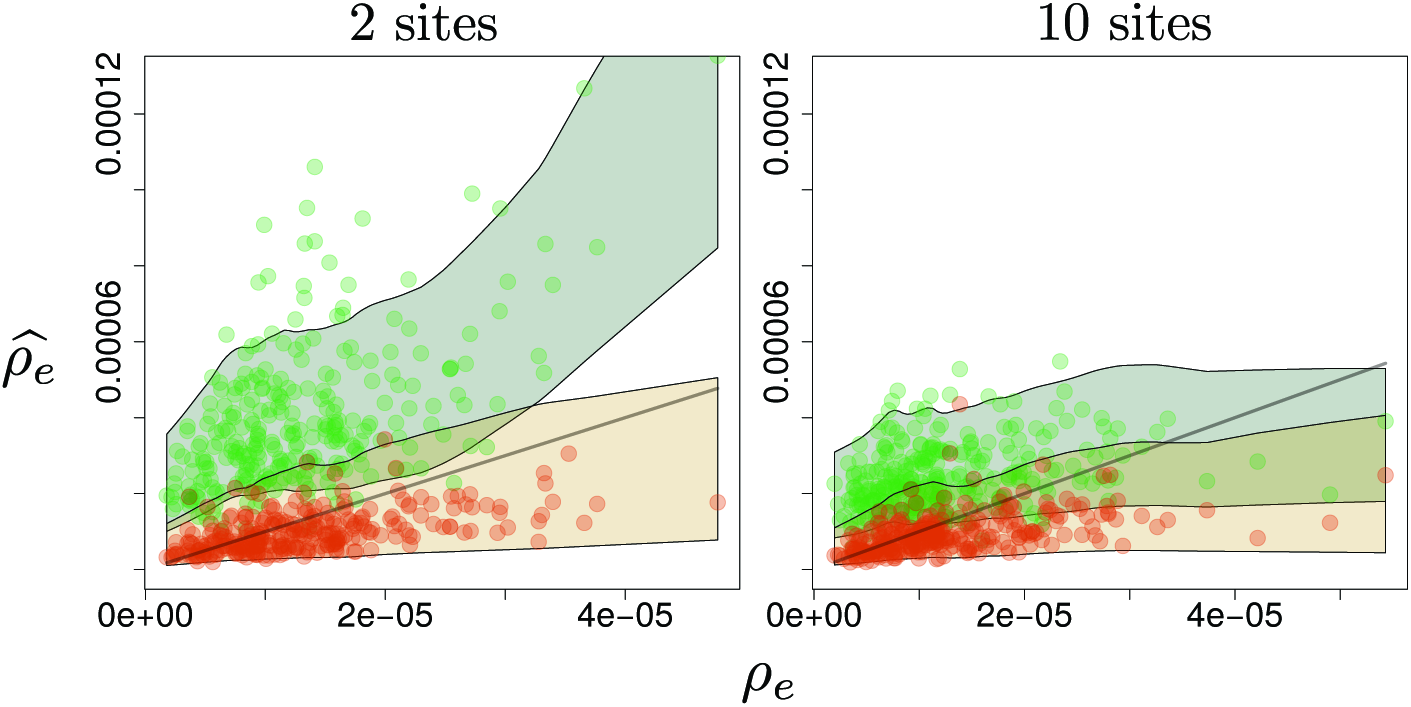
True *vs*. estimated effective population densities (*ρ*_*e*_ *vs*. 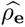) obtained under the ΛV (orange) and the structured coalescent (green) models. Left: two sampling sites. Right: ten sampling sites. 300 simulated data sets were considered in each case. The y-value for each dot corresponds to a posterior median. The upper and lower limits of the colored areas are obtained by fitting a smooth line through the 97.5% and 2.5% quantiles of the estimated posterior distributions of values of *ρ*_*e*_.

### 6-b Dispersal intensity

Figure 2 displays the simulated values of dispersal intensity (*σ*^2^) against the posterior medians inferred using the Bayesian approach. Parameter inference is fairly accurate overall but precision is limited, especially for large values of this parameter. Increasing the number of sampling regions improves the quality of inference however. Overall, these results suggest that it is possible to extract valuable information about dispersal patterns using our approach, although obtaining precise estimates requires sampling a large number of sites.

**Figure 2:**
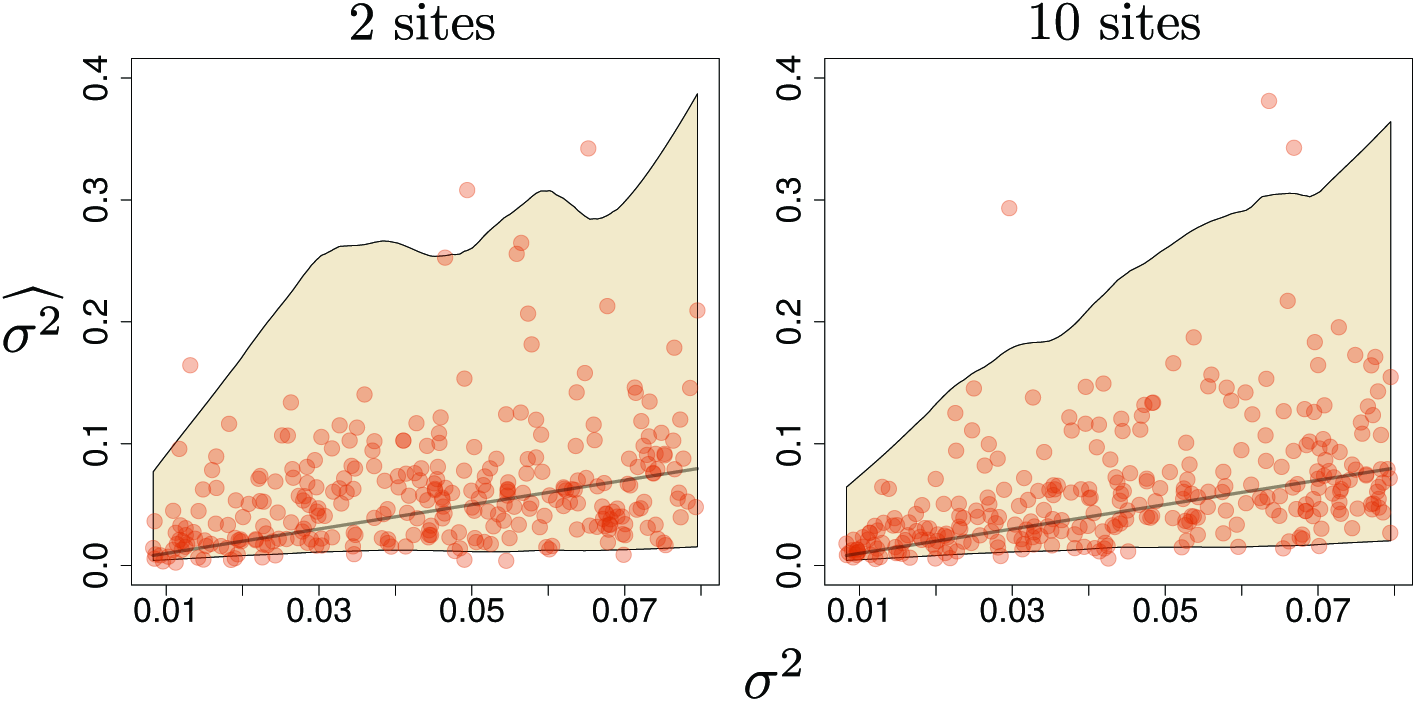
True *vs*. estimated dispersal intensity (*σ*^2^ vs. 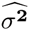) under the ΛV model. See caption of Figure 1.

### 6-c Computation times

Each simulated data set was allocated a maximum of 100 hours of computation time. Effective sample sizes (ESS) for the parameters λ, *μ* and *σ*^2^ were monitored during this period. For two sampling sites, 60%, 62% and 95% of the data sets had ESS greater than 100 for λ, *μ* and *σ*^2^ respectively. For ten sites, the corresponding percentages are 56, 80 and 99. Usable estimates of the ΛV model parameters are thus generally obtained in a reasonable, yet substantial, computation time.

### 6-d Influenza seasons in the USA

Homologous nucleotide sequences from the NA segment of the Influenza A virus (H1N1 sub-type) were retrieved from the Influenza Research Database [45]. Five flu seasons (2009-2010 to 2013-2014) in the USA were considered in five separate analyses. Hawaii and Alaska were excluded from the analysis in order to approximate the shape of the habitat with a rectangle. Multiple sequences are generally available for each state and season. Two distinct sets of sequences (with a single sequence per state for each set) were analyzed for each season, thereby providing two independent biological replicates for each of the five flu seasons. Each sequence alignment was analyzed using the HKY [19] model of nucleotide substitution with the FreeRate model of rate variation across sites [44] and a covarion-like model of site-specific rate variation across lineages [18].

Figure 3 gives the posterior distributions of the neighborhood size and the radius parameter for the five seasons and two replicates. We focused on the radius *θ* rather than the dispersal intensity parameter *σ*^2^ as the latter requires knowledge about the rate of nucleotide substitution which we could not infer because of the lack of calibration information in the data considered.

**Figure 3:**
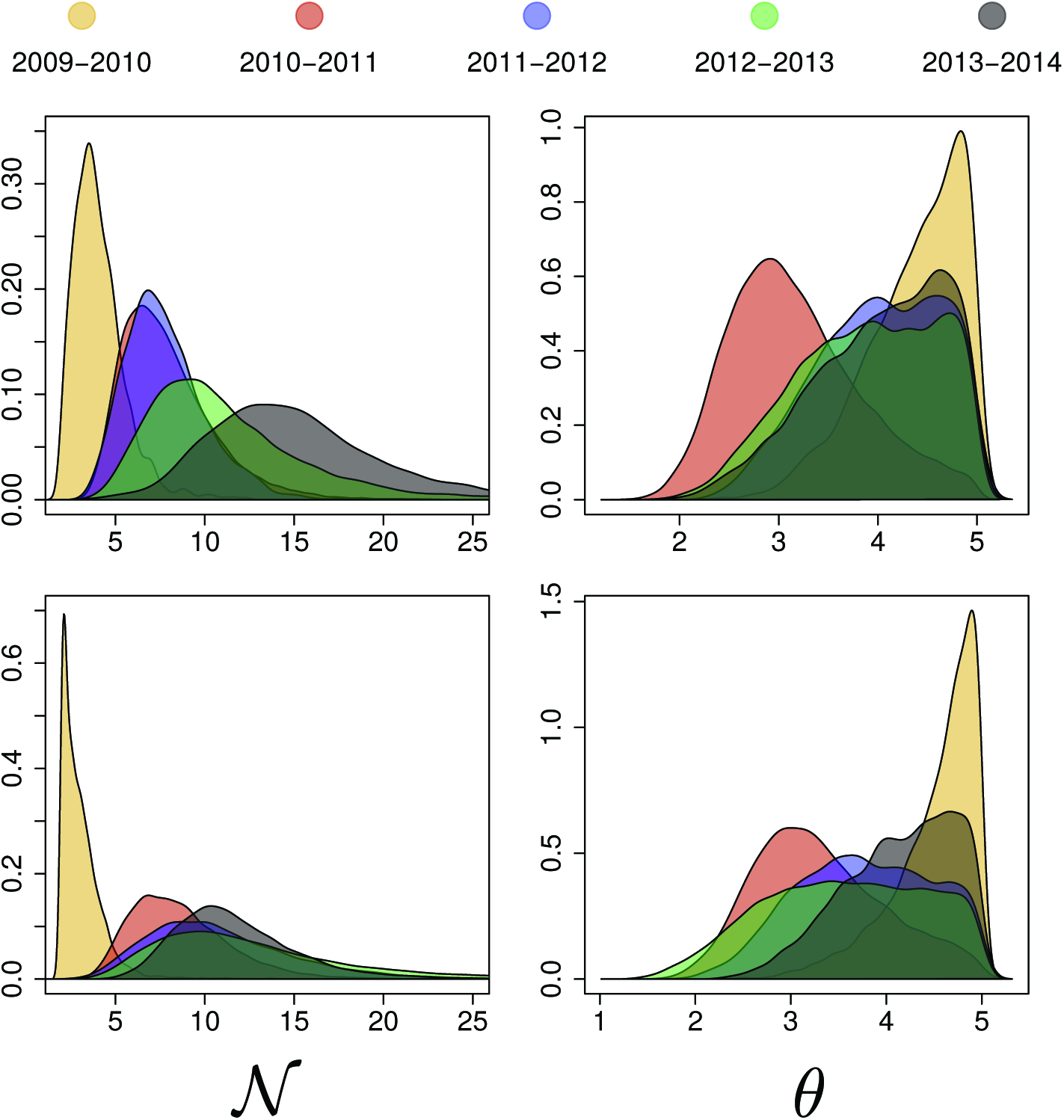
Neighborhood size (*𝒩*) and radius parameter (*θ*) estimates for five seasons of H1N1 in USA. The two rows give the results obtained with two independent biological replicates (see main text). FLU seasons are color-coded (see top).

The comparison of parameter estimates across seasons and replicates shows interesting features. First, the posterior distribution of parameters are similar in the two independent biological replicates. This observation suggests that variation of parameter estimates due to sampling is negligible. Second, the evolutionary dynamics observed for the 2009-2010 season (corresponding to the 2009 flu pandemic in the USA) are clearly distinct from that observed for other seasons. Comparatively smaller neighborhood sizes and larger radii are inferred for this season. These two observations are indicative of a virus with limited infection rate (low *𝒩*) but good ability to proliferate under various climatic conditions (large *θ*). The fact that the 2009-2010 season lasted for a substantially longer period of time compared to subsequent seasons and had a comparatively mild incidence rate (see SI 16) corroborate this conclusion.

## 7 Discussion

Understanding the forces shaping genetic diversity in space is a key objective in ecology and population genetics. Recent years have seen the rise of methods that essentially aim at visualizing the correlation between genetic and geographic distances [35, 34]. These exploratory methods can reveal interesting patterns in the data such as long-distance admixture [12], migration corridors or barriers [36]. The present study focuses instead on inferring the parameters of a stochastic model of population dynamics. This approach is well suited to testing biological hypotheses and therefore provides a relevant complement to more exploratory techniques.

Our results indicate that the spatial λ-Fleming-Viot (λV) model [14, 9, 3, 5, 47, 4] is amenable to parameter inference under biologically realistic conditions. Using a Bayesian inference technique that relies on augmenting the data with the ancestral locations of sampled lineages, we show that accurate information about neighborhood size and dispersal intensity can be recovered from geo-referenced genetic data. It is a significant step forward in the analysis of the spatial distribution of genetic diversity as partitioning populations into discrete demes (as in the structured coalescent) or assuming a non-homogeneous distribution of individuals in space and time (as in the Wright-Malécot model) is not required with this new technique.

Estimates of neighborhood sizes obtained with the traditional approach based on fixation indices (Fst) show virtually no correlation with the true values of this parameter in our simulations. This inference technique was originally designed for the analysis of diploid individuals and multiple unlinked loci with each locus evolving under an infinite-allele model. It is robust to misspecification of the mutation model [26] and is relevant in a broad range of experimental conditions [49]. In our simulations however, all loci evolved along the same genealogy while the number of alleles was limited to four nucleotides and only one sequence per individual was considered. In these circumstances, which correspond to standard experimental conditions, our Bayesian inference technique returns precise estimates of neighborhood sizes, thereby providing a relevant alternative to Fst-based methods. More importantly, while the traditional approach only estimates the product of population density and dispersal intensity (i.e., the two parameters are not identifiable in the standard inference framework, see e.g., [42]), both parameters can be estimated separately using the proposed technique.

The structured coalescent provides estimates of population density. Although this method assumes that each deme corresponds to a sub-population, it is commonplace to equate a sampling region with a deme. In these circumstances, our results suggest that the structured coalescent overestimates the population density when the generative model is λV. The bias decreases with the increase of the number of sampling regions however. Yet, our Bayesian estimation technique clearly outperforms the structured coalescent here. In cases where the population of interest is not strongly structured spatially but rather continuously distributed, we thus recommend that estimation is conducted under the ΛV model.

Bayesian inference methods are generally computationally intensive compared to other estimation techniques. Our approach is no exception, although stable estimates of the three main model parameters were obtained after about four days of computation in the majority of simulated data sets. Our implementation of the MCMC sampler fitting the ΛV model was extensively tested and optimized. Nonetheless, new operators complementing the fifteen considered here might improve the speed of convergence. Also, a substantial fraction of the computation is spent on evaluating the likelihood of the model for REX events that do not affect the location of any lineage. Integrating over the locations and times of these events could potentially be done analytically, thereby decreasing the computational burden.

The analysis of H1N1 sequences from five flu seasons in the USA provides insight into the dynamics of the infection that is coherent with the variation in the incidence of flu-related diseases. In particular, the patterns inferred for the 2009 pandemic, both in terms of neighborhood size and range of dispersal, are notably distinct from that observed in other ‘regular’ flu seasons. The larger than usual estimate of dispersal distance might be one of the factors explaining why this season lasted for a longer period of time compared to other years. Also, the incidence rate for the 2009 season was relatively mild which is consistent with the small neighborhood size estimated here.

Our implementation of the ΛV model relies on several assumptions that require careful consideration. First, the size of the population and its habitat are considered as fixed. Detecting expansion or contraction of population sizes during the course of evolution is at the core of important questions in ecology and population genetics (see e.g., [20]). Accommodating for deterministic changes of population size in the ΛV framework is therefore of utmost interest and will be the focus of future research. Second, our simulations assume a homogeneous landscape. This assumption is not realistic in instances where mountains, rivers, human activity, etc., impede migration of individuals. Relaxing the assumption of isotropic migrations in the ΛV model presents a technical challenge that needs to be addressed. Third, our simulations and inference method target multiple linked loci. Extending our approach to accommodate for recombination is a interesting research prospect. In fact, Barton et al. (2013) recently showed that recombination patterns convey information about the distribution of relative parent position and neighborhood size under the ΛV model. Fourth, the boundaries of the habitat are considered as known a priori in the present study. Treating the area of the habitat as a random variable would be relevant for the analysis of most real data sets. Further work on this question should also assess the impact of over-or under-estimating this area on the accuracy and precision of model parameter estimates.

Despite these limitations, the proposed approach is a relevant complement to the exploratory analyses mentioned above and the inferential techniques based on the structured coalescent or the fixation index. Fitting a ΛV model is particularly relevant in cases where the spatial distribution of individuals in the population of interest does not display well-defined demes. Despite the substantial computational burden involved with the Bayesian inference under this model, the opportunity to infer the density of a population and characterize dispersal distances from the combined analysis of genetic and spatial data should help improve our understanding of the mechanisms underlying spatial distribution of populations and species.

## 8 Software availability

The software phyrex implementing the MCMC algorithm for parameter inference under the ΛV model is available as part of the PhyML package from the following URL: https://github.com/stephaneguindon/phyml.

## 9 Acknowledgments

We thank Prof. Nick Barton and François Rousset for constructive feedback on early versions of the manuscript. We also wish to acknowledge the Centre for eResearch at the University of Auckland and NeSI high-performance computing facilities.

## Inference under the coalescent model in a spatial continuum

-Supplementary information-

March 2, 2016

### 10 Sampling scheme

To generate a single triangle in which individuals are sampled, three points, (*x*_*i*_, *y*_*i*_), *i* = 1, 2, 3, are first drawn uniformly at random on the square (0, *ω*/2) × (0,*h*/2). These points make up the vertices of the triangle. The resulting triangle is then translated to the right and up so that the whole triangle remains within the landscape. This is achieved by drawing *u*_1_ ~ U(max{*x*_1_, *x*_2_, *x*_3_}, *ω*) and *u*_2_ ~ U(max{*y*_1_, *y*_2_, *y*_3_}, *h*)i> and setting the vertices of the triangle to be (*x*_*i*_ + *u*_1_, *y*_*i*_ + *u*_2_), *i* =1, 2, 3. For each simulation, we iterate this procedure two or ten times to produce the corresponding number of sampling regions. The median area covered by two and ten triangles is 4% and 16% of the total area respectively, thus corresponding to a broad range of sampling conditions.

### 11 MCMC operators

We describe below the different operators, also commonly referred to as “moves”, for updating the model parameters in our Markov Chain Monte Carlo algorithm. Each operator applies a random modification to the current state of the model. These modifications and the corresponding ratios of probability densities (Hastings ratio) are detailed below.

#### 11-a Scalar scaling operator

This operator updates a scalar parameter which value is constrained to be in [*a, b*]. Let *x* and *x*′ be the current and proposed values respectively. *X* and *X*′ denote the corresponding random variables. *u* is a random draw from a uniform distribution in [0,1] (random variable *U*) and *K* is a tuning parameter. We use:

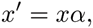

where *α* := exp{*K*(*u* −1/2)}. *α* can thus be considered as a random draw from a random variable *A* with *p*_*A*_(*α*) = 1/(*Kα*) and *e*^−*K*/2^ ≤ *α* ≤ *e*^*K*/2^. The corresponding proposal density is therefore:

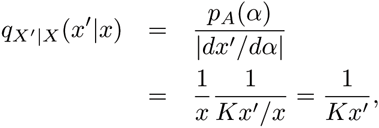

whenever *x*′ ∈ [*a, b*]. *q*_*X′|X*_^(*x′|x*)^ = 0 otherwise. Similarly, we have:

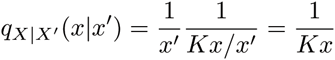

if *x* ∈ [*a, b*] and 0 otherwise. The Hastings ratio for this operator is thus:

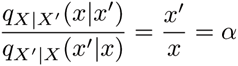

when *x* ∈ [*a, b*] and 0 otherwise. The parameters λ, *μ* and *θ*, as well as the transition/transversion ratio, are updated using this operator. The value of the tuning parameter *K* is adjusted during the first 10^6^ iterations of the Markov Chain Monte Carlo algorithm such that the frequency with which proposed values replace current ones attains 0.234 [39].

#### 11-b Vector scaling operator

Let *x*_*l*_ be a vector of non-negative real values of length *l*. 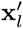j is the proposed value for that vector such that:

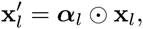

where *α*_*l*_ = (*α*_1_,…, *α*_*l*_) is the value taken by the random vector **A** = (*A*_1_,…, *A*_*l*_) and ⊙ is the element-wise product operator. The corresponding proposal density is then:

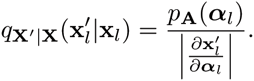

The determinant of the Jacobian is equal to 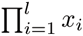. We thus re-write the proposal density:

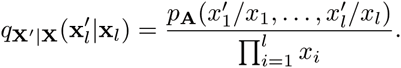

In case all values in *x* are multiplied by the same scaling factor *α*, we have 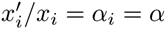 for all *i* = 1,…, *l* and we can write:

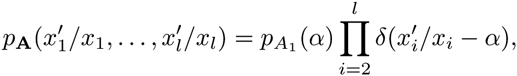

where *δ*(.) is the Dirac delta function. We thus have:

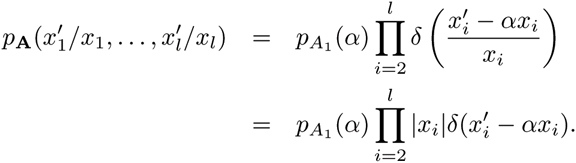

Similarly, we have:

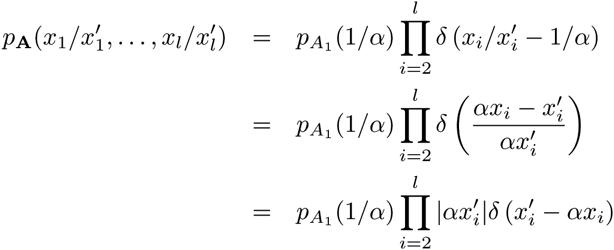

Therefore,

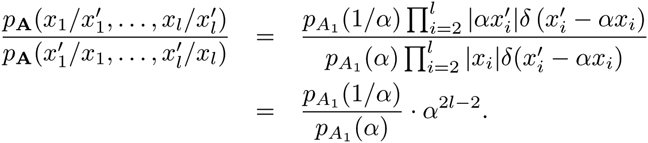

The Hastings ratio is therefore obtained as follows:

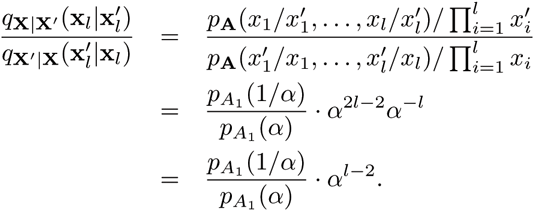

Using *p*_*A*_1__(*α*) = 1/(*K ˙ α*) with *e*^*−K/2*^ ≤ *α* ≤ *e*^*K*/2^ as for the scalar scaling operator, we obtain:

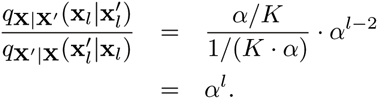

The vector scaling operator is used for scaling the times of REX events up and down. This operator is used in conjunction with another operator that updates the value of λ. Indeed, the timing of REX events results from a Poisson process with rate λ. It therefore makes sense to update these parameters together. In particular, let *T*_*h*_ and 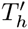 = *αT*_*h*_ be the current and proposed time of the MRCA of the sampled lineages (i.e., the height of the tree) and *m* the total number of REX events. A new value of λ is obtained by dividing the current value of this parameter by *α*. The Hastings ratio for this sub-operator is 1/*α*.

#### 11-c Insert/delete REX events without lineage displacement

This operator adds or removes multiple REXs that do not affect the spatial coordinates of any of the sampled lineages. Each of these events takes place at a time between present (*t*_0_ = 0) and the time of the MRCA for the sampled lineages (*T*_*h*_ = *t*_*h*_^1^). Let *m* and *m*′ be the current and proposed number of these events. *M* and *M′* are the corresponding random variables. *M′* follows a Poisson distribution with parameter m. The proposal distributions are thus:

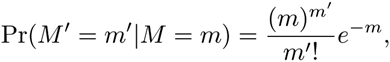

and

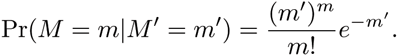

The Hastings ratio for that part of the operator is derived from these two probabilities. Once *m′* is known, one needs to set the times of newly inserted events or remove some of the existing ones. We distinguish two cases: *m′* > *m* and *m′* < *m* (if *m′* = *m*, the model remains unchanged). If the first case, *n*_+_ = *m′* − *m* REX new events are inserted. The times at which these events take place are chosen uniformly at random in [*t*_*h*_, *t*_0_]. The corresponding joint density of the new times is thus (*n*_+_)!. (1/|*t*_*h*_|)^n+^. In the second case, *n*_−_ = *m* − *m′* existing REX events are removed from the model. The probability of removing a particular set of *n*_−_ events in a model with currently *m* such events is 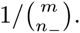. The Hastings ratio corresponding to the insertion of *n*_+_ events is thus:

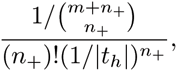

and the Hastings ratio when removing *n*_−_ events is as follows:

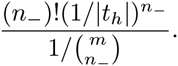

The third and last sub-operator concerns the spatial coordinates of the centers for each inserted REX event. These coordinates are sampled uniformly at random on the landscape of surface area *s*. When inserting *n*_+_ new REXs, the Hastings ratio for this part of the operator is thus:

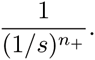

When deleting *n*_−_ REXs, the Hastings ratio is simply:

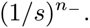

In summary, this operator is divided into three sub-operators. The Hastings ratios corresponding to these three sub-operators are multiplied together to determine the Hastings ratio for the whole operator. As for the operator that rescales the tree height, the value of λ is also updated in the insert/delete operator since the number of REX events taking place in a given time period is highly correlated to that of λ. A new value for this parameter is proposed by sampling from a left-and right-truncated normal distribution with mode equal to *m′/t*_*h*_ and standard deviation equal to 0.1 *m′/t*_*h*_. The normal distribution is left-truncated at 0.0 and right-truncated at 1.2 .*m′/t*_*h*_ (i.e., the mode plus two times the standard deviation). We used the accept-reject algorithm described in [38] to sample from univariate truncated normal distributions.

**Figure 4:**
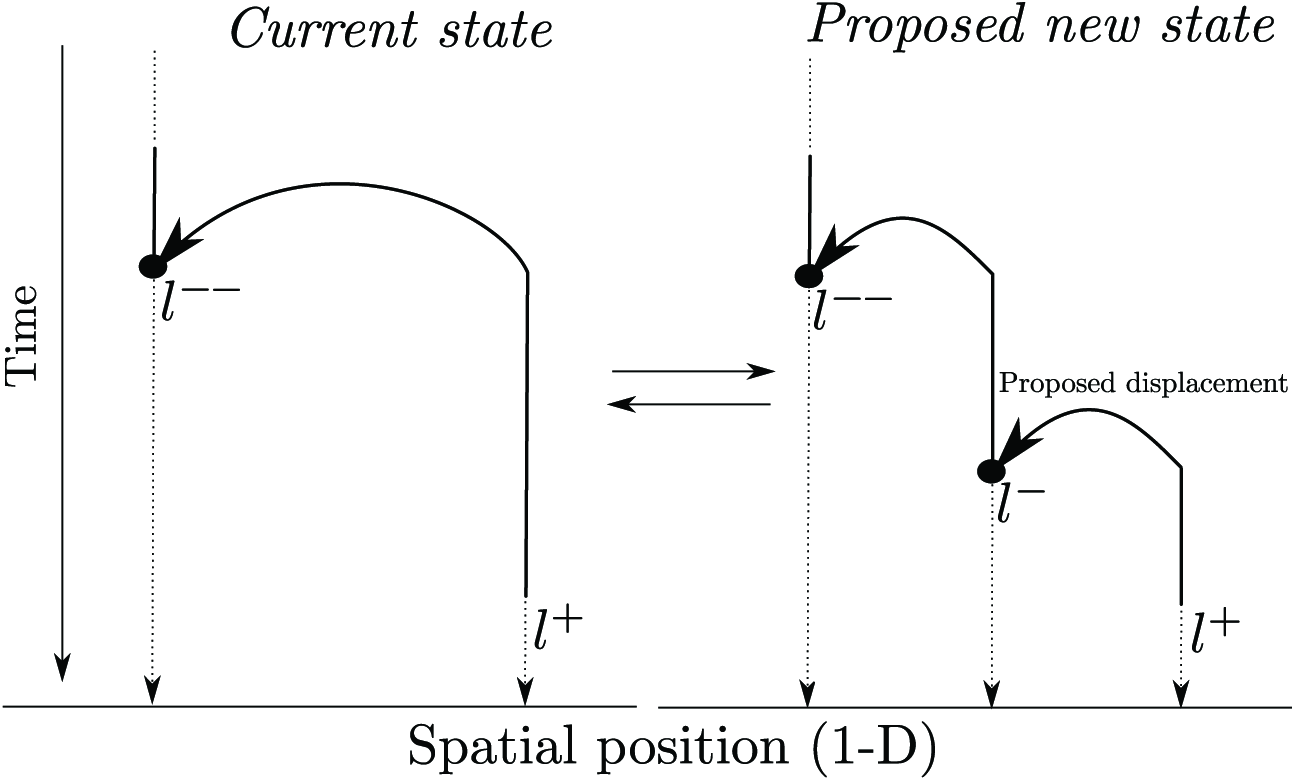
Addition (and removal) of a REX event with lineage displacement. *l*^+^ and *l*^−^ are the coordinates of the lineage (projected on a single axis) immediately before and after the proposed new REX event respectively. *l*^−−^ is the position of the same lineage immediately after the last REX event (after the proposed one) that has hit this lineage.

#### 11-d Insert/delete REX events with lineage displacement

This operator adds or removes one or more REXs, each of them hitting a sampled lineage (i.e., REXs corresponding to coalescent events and those leaving the spatial position of all sampled lineages unchanged are not affected by this operator). The three sub-operators described in the previous section also apply here. An additional sub-operator deals with the spatial coordinates of the newly proposed REXs in case 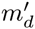 > *m*_*d*_, where 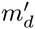 and *m*_*d*_ are the proposed and current number of REX events with lineage displacement in the tree. Let *L*_+_ be the random variable corresponding to the new spatial coordinates of one of the lineages hit by one of the *n*_+_ proposed REX events (with *n*_+_ = 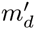 − *m*_*d*_ > 0). *C*_+_ is the random variable corresponding to the coordinates of the center of this REX and *c* is the value taken by *C*_+_. *l*^−^ is the value taken by *L*_+_, while *l*^+^ is the coordinate of the same lineage just before the displacement (going backward in time). *l*^−^ is the position of the same lineage immediately after the next REX event that affected it (see Figure SI 4).

We take *L*_+_ as a normally distributed random variable with mode (*l*^−^ + *l*^+^)/2 and variance 2*θ*^2^. This distribution is left-and right-truncated (at 0 on the left and *h* or *ω* of the right) in order to avoid having lineages migrating beyond the limits of the habitat. Let *p*_*L*_+__ (*l*^−^|*l*^+^, *l*^−^, *θ*) denote the corresponding density. The distribution of *C*_+_ is also a left-and right-truncated normal. The mode of this distribution is *l*^−^ and the variance *θ*. Let *p*_*c*_+__(*c|l*^−^, *θ*) denote the corresponding probability density. Altogether, the joint probability density for the proposed lineage and center positions is as follows:

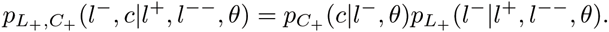

Since each REX event with a lineage displacement can be chosen for removal with equal probability, the Hastings ratio for inserting one such event is thus:

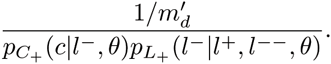

The Hastings ratio for deleting one event is as follows:

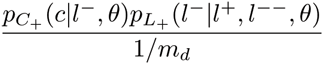

When adding or removing *k* ≥ 1 REX events with lineage displacements, the Hastings ratio are:

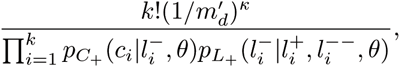

for the insertions and

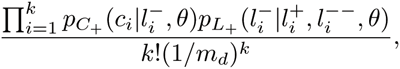

for the deletions. As for the insert/delete REX without displacement operator, the number of REX events inserted or deleted, *k*, is a random draw from a Poisson distribution. The parameter of this distribution is given by the number of REXs with lineage displacement in the current tree (or the proposed tree for the reverse modification that updates the proposed model into the current one). The corresponding two probabilities define the Hastings ratio for this sub-operator.

#### 11-e Path operator

This operator updates the series of displacements between a coalescent node (the father) and one of the coalescent node immediately below the first (the daughter). Let *l*_*f*_ and *l*_*d*_ be the coordinates of the lineage immediately after the father and the daughter nodes arise in the genealogy. Let *ω* be the current number of lineage displacements between the two coalescent nodes and*ω′* the proposed new number of displacements. (*l*_*i*_,…, *l*_*ω*_) and (*l′_1_,…*, *l*′_*ω*_) are the current and proposed coordinates of the lineage under study. Figure SI 5 gives an example illustrating this operator.

**Figure 5:**
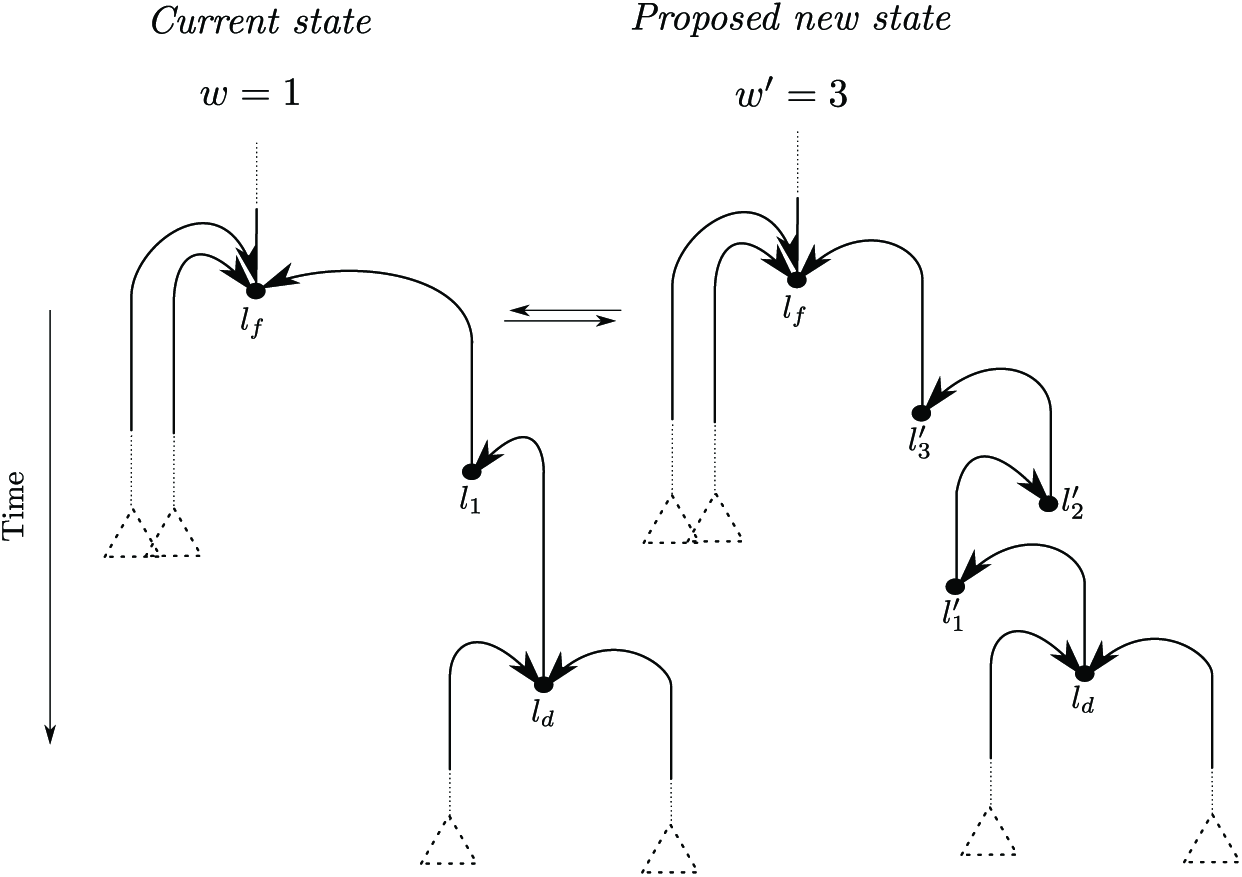
Path operator. *l*_*f*_ and *l*_*d*_ are the locations of the lineage under scrutiny immediately after the corresponding REX events. In the current state of the model, a single lineage displacement occurs between the father and daughter coalescent nodes, thus *ω* = 1 and *l*_1_ is the position of the lineage immediately after the displacement occurs. In the proposed new state, *ω′* = 3 and the locations of the lineage just after the corresponding REXs are *l′*_1_, *l*′_2_ and *l*′_3_.

The proposed number of lineage displacements *ω′* is a random draw from a Poisson distribution with parameter obtained by counting the current number of displacements in the tree (excluding coalescent events) and dividing by the sum of edge lengths (in calendar time unit). The probability of having *ω′* displacement(s) between the father and daughter nodes is thus

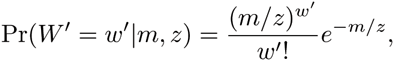

where *m* is the current total number of lineage displacements in the genealogy, excluding coalescent events and *z* is the sum of edge lengths.

The conditional distribution of 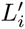 given 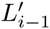(with *i* = 1,…, *ω′* and 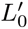 = *l*_*d*_) is a left-and right-truncated normal with variance 2*θ*. The mode of that distribution is 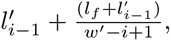>, with 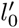 = *l*_*d*_. The rationale behind shifting the mode of the normal distributions is to implement a sampling strategy similar to that used for generating a Brownian bridge. Lastly, the conditional distribution of the center 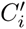 given 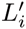 is a left-and right-truncated normal with mode set to the value taken by 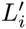 and variance *θ*.

Altogether, the proposal density for this operator is then:

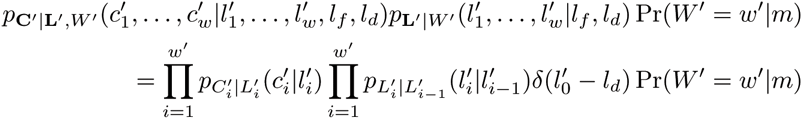

where 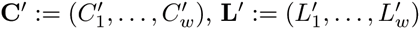 and **L** := (*L*_1_, …, *L*_*ω*_ The very same reasoning is used to derive the proposal density for the reverse modification of the model and the Hastings ratio for the path operator follows.

#### 11-f Subtree Prune and Regraft (SPR) operator

This operator changes the topology of the tree. A coalescent node is first sampled uniformly at random. The root node is included in the list of coalescent nodes that can be chosen from only if its degree is strictly greater than two. Let *n*_*p*_ be the number of “valid” coalescent nodes in the current tree. The coalescent node selected has *n*_*p,d*_ direct descendants, where each direct descendant is the first (coalescent or tip) node encountered when traversing the tree starting from the selected node along each of the *n_p_d* paths leading to a tip. In the example given in Figure SI 6, the selected coalescent node is noted as *𝒫* and *n*_*p,d*_ = 3. One of the *n*_*p,d*_ descendant nodes is chosen uniformly at random (noted as *𝒟* in Figure SI 6). The subtree with the selected descendant node as its crown is pruned. The position in the tree at which the pruned subtree is re-grafted can be either a coalescent node or, as in Figure SI 6, a “displacement node”, i.e., a node of degree two corresponding to the displacement of a lineage. The re-graft node is noted as *𝒢*. The corresponding time for that node has to be older than that of *𝒟*. Also, the re-graft node cannot be on the path from *𝒫* to *𝒟*. Let *n*_*g*_ be the number of valid re-graft nodes. The pruned subtree is re-grafted to the tree after generating a new series of lineage displacements between *𝒟* and *𝒢* using the path operator.

**Figure 6:**
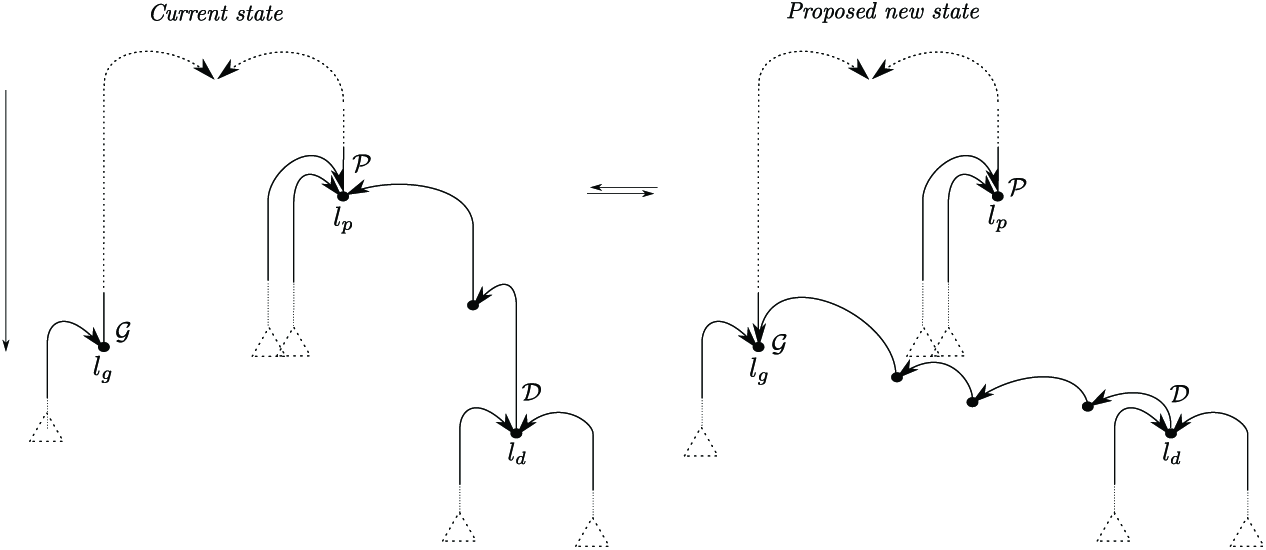
SPR operator. *𝒫, 𝒟* and *𝒢* are the selected nodes for pruning, its selected descendent and the re-graft node respectively.

The joint probability of selecting nodes *𝒫* and *𝒟* is equal to (1/*n*_*p*_)(1/*n*_*p,d*_). The probability of selecting *𝒢* as re-graft node is 1/*n*_*g*_. Altogether, the probability of pruning a particular subtree and re-grafting it is thus (1 /*n*_*p*_)(1/*n*_*p,d*_)(1/*n*_*g*_). The probability of reverting this topological change is obtained after updating the values of *n*_*p*_, *n*_*p,d*_ and *n*_*g*_ on the proposed new tree. The Hastings ratio for the SPR operator also involves the ratio of densities of the (new and exisiting) paths from *𝒟* to *𝒫* and *𝒟* to *𝒢* (see previous section).

#### 11-g Backward simulation operator

This operator relies on the backward-in-time dynamics of the ΛV model. A time point, *t*, is first chosen uniformly at random in the [0,*t*_*h*_] interval. Let *t*_*i*_ be the most recent REX that occured after *t*. All the REXs strictly older than ti are then discarded. A new “upper” part of the genealogy is then simulated according to the ΛV model, using the current values of λ, *μ* and *ρ*. The Hastings ratio for this operator is then as follows:

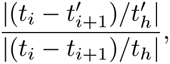

where 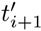 is the proposed time of the most recent REX that occured before *t* and 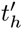 is the time for the MRCA in the proposed genealogy.

#### 11-h Other operators

Additional operators were used in our implementation of the MCMC sampler for estimating the parameters of the ΛV model. These extra operators are more straightforward than those described above, with Hastings ratios that obtained immediately. For this reason, we do not give a detailed mathematical description of these operators.

The first operator changes the time of a REX event with lineage displacement (which can also be a coalescent node). The proposed new time is sampled uniformly at random in the interval with upper and lower bounds corresponding to the next and previous REX events in the current model respectively. Note that the two REXs defining the upper and lower bounds involve lineage displacements or not. This operator is applied simulateously to a fixed fraction of the current number of REX events in the model. Its Hastings ratio is equal to one. A similar operator is also implemented whereby the proposed time of a REX with lineage displacement is uniform between the next and previous REXs with lineage displacement. This operator is applied to single REX events. Its Hastings ratio is also equal to one.

Beside operators that update the time of REXs, we also found it helpful to implement operators that potentially modify the spatial coordinates of lineages and REX centers. In the first case, the coordinates of a lineage immediately before a REX with lineage displacement are sampled from two truncated normal distributions independently (one draw for the lattitude, one for the longitude). The mode of each truncated normal is given by the centre of the corresponding REX event. The standard deviation is set to *θ*, the current value of the radius parameter. This operator was applied simultaneously to a fixed fraction of REXs with lineage displacement. For each REX, the Hastings ratio is determined by the densities of the aforementionned truncated normal distribution evaluated at the new and the current lineage coordinates. A very similar operator is applied to the centers of REX events with lineage displacements. The modes of the truncated normal distributions are determined here by the coordinates of the corresponding lineages immediately before the REX events involved.

### 12 Interpretation of parameters

When considering the backward-in-time process, the rate at which a lineage is hit by a REX is the product of the rate at which these events occur (λ) by the probability that a lineage is hit. Let 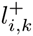 be the location of lineage *k* just before the REX event that occured at time *t*_*i*_ (going backward in time). The probability that this lineage is hit conditional on the REX having location *c*_*i*_ := (*c*_*x*_, *c*_*y*_) is

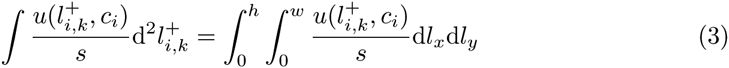

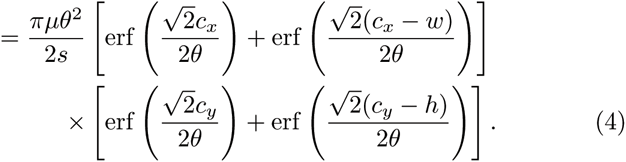

In cases where the argument of the error functions is large enough (i.e., greater than ⋍ 2), the value returned is approximately equal to one. These conditions are met when *θ* ≪ min(*c*_*x*_, *c*_*y*_) and *c*_*i*_ is far enough from the edges of the habitat (i.e., *c*_*x*_ ≪ *ω* and *c*_*y*_ ≪ *h*). In this situation, the expression above simplifies and gives

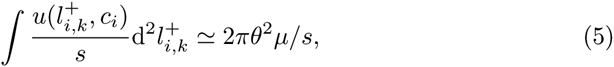

which is also the unconditional probability of the lineage being hit. We will consider that this approximation holds in what follows. The rate at which a given lineage is hit is thus 2*λπθ*^2^*μ/s*. Also, the probability density of 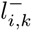 given 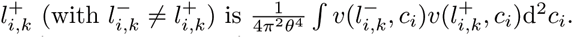 This integral yields 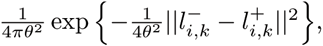 i.e., a bivariate normal density with mean 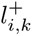 and covariance matrix 2*θ*^2^**I**. The variance of offspring location (noted as 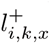) in a one-dimensional space given the parental location 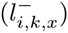 is thus:

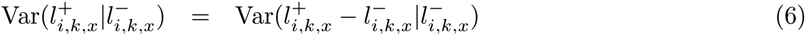

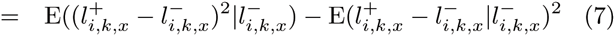

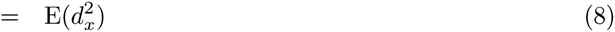

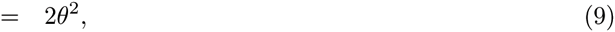

where 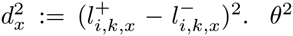 is thus half the expected square Euclidean distance between parent and offspring in one dimension. Note that 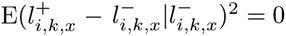 since migrations are isotropic. In two dimensions, we have:

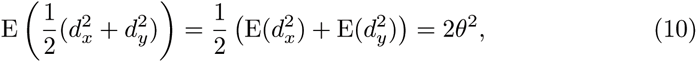

i.e., *θ*^2^ is thus a quarter of the expected square Euclidean distance between parent and offspring. In a *n*-dimensional space, *θ*^2^ is 1/2*n* times this expected distance.

Altogether, the variance of spatial coordinates of a lineage along a given axis thus increases with time proportionally to

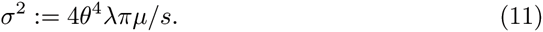

In the limit where λ → ∞ and θ → 0, the backward-in-time motion of a single lineage is a Brownian process with diffusion parameter *σ*^2^.

Also, given REX *i* occurs at position *c*_*i*_ := (*c*_*x*_, *c*_*y*_), two lineages *j* and *k* with locations 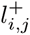 := (*l*_*j,x*_, *l*_*j,y*_)and 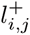 := (*l*_*k,x*_, *l*_*k,y*_), in a slight simplification of the notation used previously, coalesce with probability

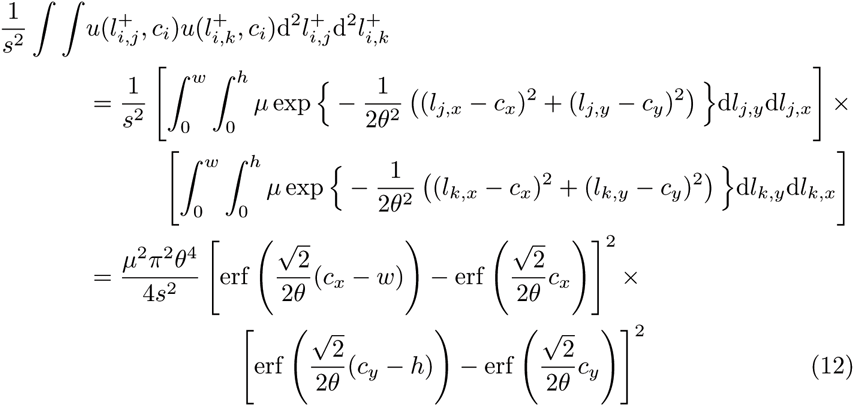

In cases where the radius is small (see above), the probability that the two lineages coalescence, conditional on the position of the REX event is thus approximately equal to 4*μ*^2^π^2^θ^4^/s^2^, which also corresponds to the probability of coalescence of two lineages without conditionning on the position of the REX. Note that this probability could have been derived immmediately by noting that, conditional on the location of the REX event, the event “lineage *j* is hit” is independent from the event “lineage *k* is hit”. The probability of interest is thus simply the product of the probability of each lineage being hit in the event. The rate at which pairs of lineages coalesce is therefore equal to 4*μ*^2^π^2^θ^4^λ/s^2^, which we define as 1/(2*N*_*e*_), where *N*_*e*_ is the effective population size. We therefore have

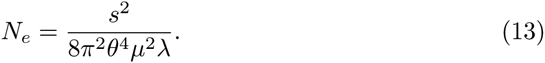

We now define the neighborhood size as *𝒩* := 4*πN*_*e*_σ^2^/*s*, following the definition used in the Wright-Malecot model. Replacing *σ*^2^ by the expression given in Equation 11 and *N*_*e*_ by that in Equation 13 yields *𝒩* = 2/*μ*.

### 13 Validation of the implementation

We validated our implementation of the simulation and inference techniques by sampling from the distribution of the following random variable

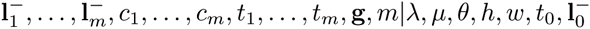

in two different ways. Values of λ, *μ* and *θ* were fixed to the median values used in our simulations throughout. Also, the locations observed at the tips of the tree (i.e., values of 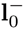) were equally spaced on the habitat. The first sampling technique relies on the functions used to simulate data backward-in-time under the ΛV model. The second technique relies on our MCMC sampler, which was slightly modified such that values of λ, *μ* and *θ* were fixed to the median values aforementionned. 3,000 data sets were generated using the simulation technique while the Bayesian sampler was run for approximately 10 hours.

**Figure 7:**
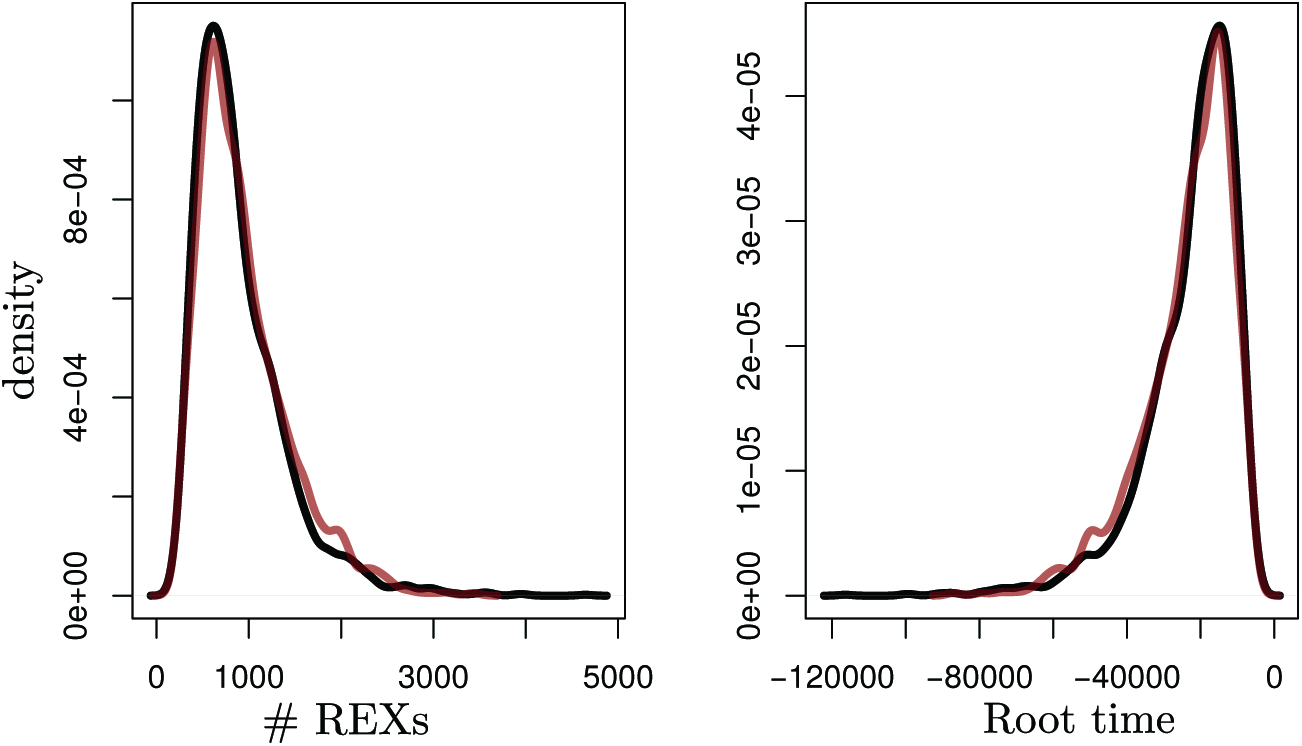
Distribution of the number of REX events and the tree height obtained with our simulation and MCMC sampling techniques. Black: distributions obtained with the backward-in-time simulation-based approach. Purple: distributions obtained with our MCMC sampler.

Figure SI 7 shows the marginal distributions of two summary statistics: the number of REX events (i.e., *m*) and the time (before present) at the root of the tree (i.e., *t*_*m*_). The distributions obtained with both sampling techniques are virtually identical, suggesting that our implementation of the backward-in-time simulation of the ΛV model is consistent with the corresponding distribution sampled by our MCMC.

### 14 Estimation of neighborhood size using fixation index

The relationship between logarithm of spatial distance (*r*) and fixation index (*F(r)*) for pairs of sampled individuals is linear in two-dimensional landscapes [41, 48, 2]. The slope of the corresponding linear regression, noted as *α*, then serves as a basis for the inference of Wright’s “neighborhood size”. Indeed, assuming a Gaussian dispersal function, the equality *α* = (*F*(0) − 1)*s*/4*πN*_*e*_*σ*^2^, i.e., *α* = (*F*(0) − 1)/*𝒩*, holds at equilibrium.

Values of *F(r)* were estimated for each pair of sequence with the ratio 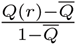, where *Q(r)* is the proportion of sites with identical states when the two sequences in the pair are at distance *r*. 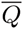 is the mean of *Q(r)* values taken over all the distinct pairs of sequences. Estimates of *F*(0) were obtained for each data set by taking the average of the fixation index values for every individual and its closest neighbor, as suggested by [48].

### 15 Bayesian estimates of neighborhood sizes

Figure 8 shows the scatterplot of simulated *vs*. estimated values of the neighborhood size (*𝒩*) obtained under the ΛV model.

**Figure 8:**
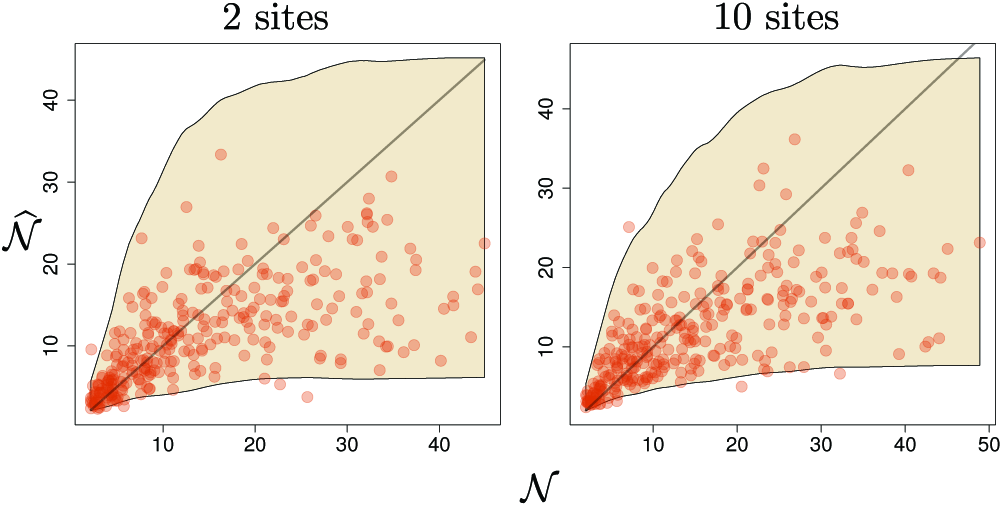
True *vs*. estimated values of the neighborhood size (*𝒩* vs.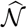) For each number of sampling sites (two on the left and ten on the right), 300 simulated data sets were analysed. The *y-*value for each dot corresponds to the posterior median estimate for a given data set. The upper and lower limits of the colored areas are obtained by fitting a smooth line through the 97.5% and 2.5% quantiles of the posterior distributions of values of *𝒩*.

### 16 Laboratory-confirmed influenza hospitalizations

Information about the incidence of influenza in the USA was gathered from the Centre for Disease Control and Prevention web site. Cumulative incidence rates per 100,000 population in the USA were fetched from the Influenza Surveillance Network (FluSurv-NET) database. FluSurv-Net covers over 70 counties in more than ten states: CA, CO, CT, GA, MD, MN, NM, NY, OR and TN were sampled every flu season. IA, ID, MI, OK and SD were also sampled in the 2009-2010 season; ID, MI, OH, OK, RI and UT were sampled in the 2010-2011 season; MI, OH, RI and UT during the 2011-2012 season, IA, MI, OH, RI and UT during the 2012-2013 season; and MI, OH and UT during the 2013-2014 season.

**Figure 9:**
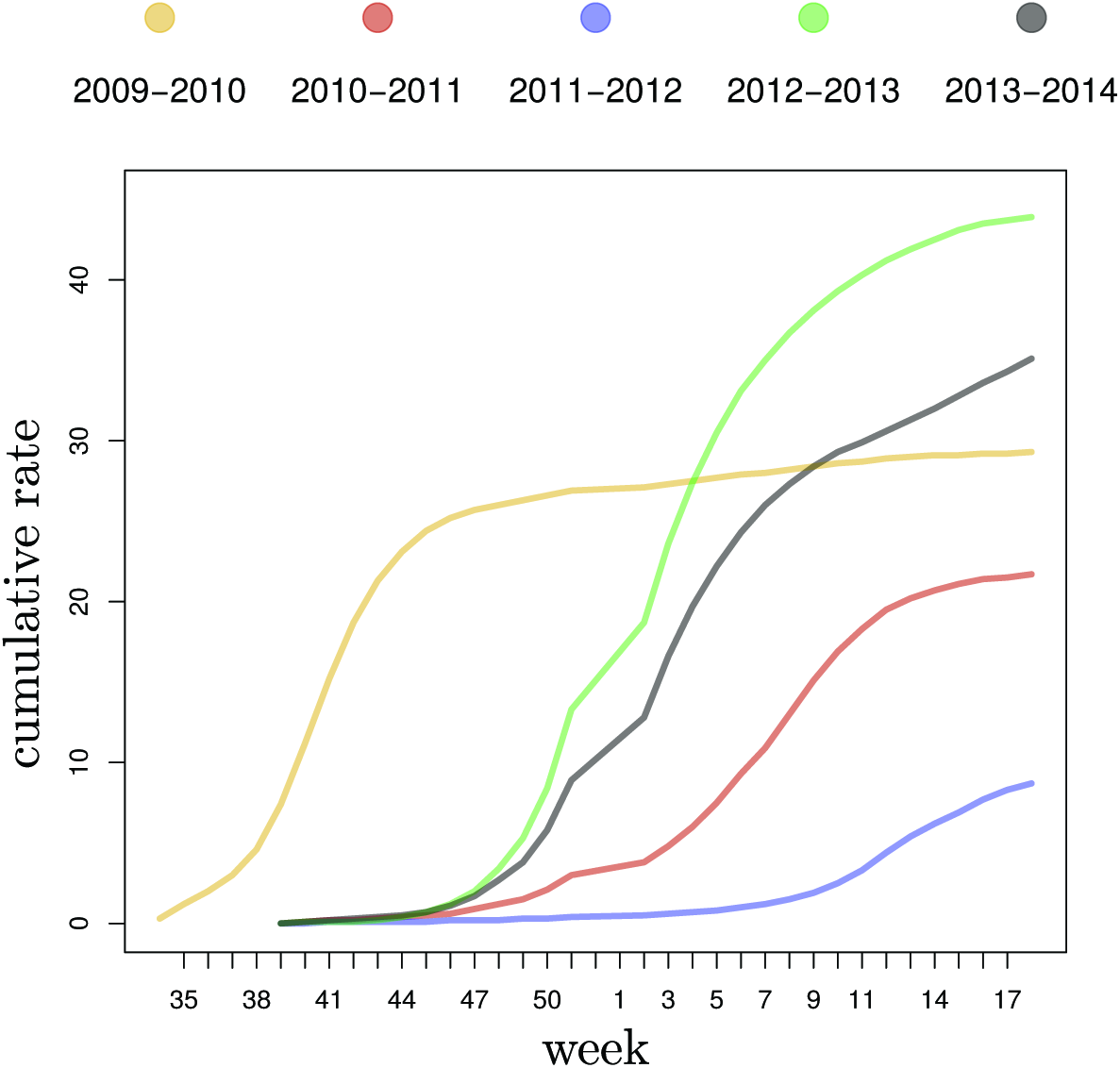
Cumulative incidence rate of individuals (per 100,000 population in the USA) that underwent laboratory-confirmed influenza related hospitalization. Data from http://gis.cdc.gov/GRASP/Fluview/FluHospRates.html.

### 17 Estimation of coalescence rates using Multi-TypeTree

The MultiTypeTree software from the BEAST2 software package [46, 11] was used to estimate the rate at which pairs of lineages coalesce under the structured coalescent model in our simulations. The number of demes was set to two or ten depending on the simulation setting. Relative rates of migration between demes were estimated from the data. Log-normal prior distributions with default hyper-parameter values were used here. The rate of coalescence was estimated in each deme and a log-normal distribution with default hyper-parameter values was also used to model the prior distribution of the inverse of each of these rates. Nucleotide sequence alignments were analysed under an HKY model [19] with a log-normal prior distribution of the transition/transversion ratio. This ratio along with the nucleotide posterior frequencies were estimated from the data. The substitution rate was fixed to 1.0 expected substitution per site per calendar time unit thoughout the tree (i.e., we assumed a strict-clock model). We used the default values for the tuning parameters and weights of the various MCMC operators available in BEAST2.

The structured coalescent permits the estimation of the rate at which pairs of lineages coalesce in a set of demes. Let *r*_*i*_ denote the rate of coalescence in deme *i* and *K* the number of demes. Considering all demes, the rate at which coalescence events occur is 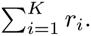. The effective size of a diploid population can be defined as 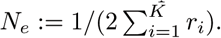. The effective population density is then *ρ*_*e*_ := *N*_*e*_/*s*. The 2.5%, 50% and 97.5% estimates of the posterior quantiles of *ρ*_*e*_ were obtained from the values of *r*_*i*_ in each deme sampled from their posterior distributions and given as output of MultiTypeTree.

## Footnotes

1 This notation slightly deviates from the one used on the main text where the time of the MRCA is noted as *t*_*m*_.

